# Anticipated distractor processing relies on distinct representational states before and after distractor onset in visual working memory

**DOI:** 10.64898/2026.09.14.751436

**Authors:** Kexin Wang, Carol A. Seger, Qi Chen, Canhuang Luo

**Author notes:** Corresponding author, (Q.C.); (C.L.).

## Abstract

Successful goal-directed behavior requires limiting the impact of salient information that is irrelevant to current goals. When distractors can be anticipated, task-irrelevant information may be encoded before it appears, yet the functional role of such anticipatory coding remains unclear. Does advance coding of irrelevant content itself support protection of visual working memory (VWM), or does effective control depend on how distractor representations evolve after entering processing? Here, using human electroencephalography (EEG) and multivariate decoding, we addressed this question by tracking distractor representations across temporal, spectral, and spatial dimensions. The distractor category was cued in advance and reliably decoded before physical onset, demonstrating content-specific anticipatory coding. However, the strength of this coding was not associated with behavioral protection. Instead, distractor processing showed a structured transition from relatively stable and temporally generalizable pre-distractor coding to more temporally unstable and time-specific post-distractor coding. Feature-level analyses further showed that distractor representations were shifted into a different representational configuration relative to representations of the same features when they served as task-relevant targets, indicating post-distractor reformatting of irrelevant content. Spectral and spatial analyses further showed that pre-and post-distractor coding relied on distinct frequency and scalp-topographic signatures. Strong anticipatory coding was associated with reduced subsequent post-distractor instability, whereas behavioral protection was associated with post-distractor instability even when pre-distractor metrics were taken into account. Together, these findings suggest a temporally coordinated control mechanism in which anticipatory coding may specify the upcoming irrelevant content, whereas post-onset reformatting may implement control by placing that content into a less stable representational state to reduce behavioral interference.

## Introduction

Goal-directed behavior requires the brain to prioritize information that is relevant to current goals while limiting the impact of salient but irrelevant input. This challenge becomes more complex when distraction can be anticipated: advance information about upcoming distractors may support preparation, but it also raises a paradox—effective control may require representing information that should ultimately be ignored. Thus, if task-irrelevant content is represented before it appears, what function does this anticipatory coding serve? Does anticipatory coding itself constitute a protective state, or does its functional role depend on how distractor representations are subsequently processed?

A large body of research has shown that the brain can use advance information to reduce the impact of distraction. Distractor suppression can be guided by spatial cues, feature cues, temporal expectations, or learned regularities, and such anticipatory states have been associated with improved selection and task performance [14, 29, 33–35]. Work on distractor templates and templates for rejection further suggests that the brain can use information about “what to ignore” to guide selection away from irrelevant input [15, 33]. In visual working memory (VWM), such anticipatory preparation has also been linked to mechanisms that protect maintained representations from upcoming interference, including alpha-band activity associated with sensory gating [3, 22, 40]. Together, these findings establish that the brain can prepare for upcoming distraction and that such preparation can influence subsequent processing. However, they do not address what role the representation of the anticipated distractor category may play in preparation. A detectable pre-distractor representation could reflect an effective template for limiting interference, but it could also serve a preparatory function that shapes how distractor representations are subsequently processed.

A detectable pre-distractor representation can show that the brain has anticipated what may appear, but it cannot by itself reveal how the distractor will be managed once it appears and competes with maintained information. In models of attention and working memory, traditional accounts often emphasize active gating or gain reduction to weaken incoming irrelevant input [3, 23], whereas complementary representational views suggest that distraction may also be managed by transforming, deprioritizing, or separating irrelevant information from task-relevant representations [2, 10, 28]. Regardless of whether control is expressed through weakened input processing or altered representational format, both perspectives highlight the need to look beyond the pre-stimulus period. Understanding the functional role of anticipation therefore requires tracking how distractor representations are organized over time: whether pre-and post-onset coding reflect a continuous representational state, or whether they are better characterized as distinct states with different functional relationships to behavioral protection.

The present study asked the question of what function anticipatory distractor coding serves in VWM. We used a delay-period distractor paradigm in which the category of upcoming task-irrelevant stimuli was cued and therefore could be anticipated, allowing us to track distractor representations before and after their physical onset. An angle detection (AD) task allowed us to test whether distractors preserved the representational format observed when the same stimuli were task-relevant, or formed a different representational configuration after onset. Using human electroencephalography (EEG) and multivariate decoding, together with temporal generalization, frequency-resolved analyses, and spatial weight patterns, we tested whether anticipatory distractor coding forms part of a continuous protective state or whether protection is better explained by the post-onset dynamics of distractor representations. We found that task-irrelevant categories were encoded before their physical onset, but this anticipatory coding did not by itself account for protection of VWM. Instead, pre-and post-onset states were distinct yet statistically coordinated, and behavioral protection was linked specifically to post-onset dynamics. These findings suggest a temporally structured and coordinated control process in which anticipatory coding reflects an early preparatory state, whereas post-onset representational reorganization may constitute a key process through which interference is controlled.

## Results

We recorded EEG data from 30 healthy participants while they performed a VWM task and an AD task (Fig 1A, B). The task-irrelevant distractor category (grating, moving dots, or line stimuli) was cued at the beginning of blocks of trials in each task (Fig 1C); this pre-cueing design allowed us to track distractor representations before and after distractor onset. In the VWM task, on each trial participants memorized the orientations of two bars and recalled the orientation of the cued bar after a delay period. During the delay, task-irrelevant distractors (grating, moving dots, or line stimuli) were presented. In the AD task, the same stimulus categories served as task-relevant targets, and participants judged for each stimulus whether its orientation/direction was horizontal or vertical. This task enabled comparison of representations under different task contexts. We first examined whether anticipated distractor categories could be decoded before onset, then characterized how distractor representations evolved after onset and whether these dynamics were associated with behavioral protection against interference.

**Fig 1.**
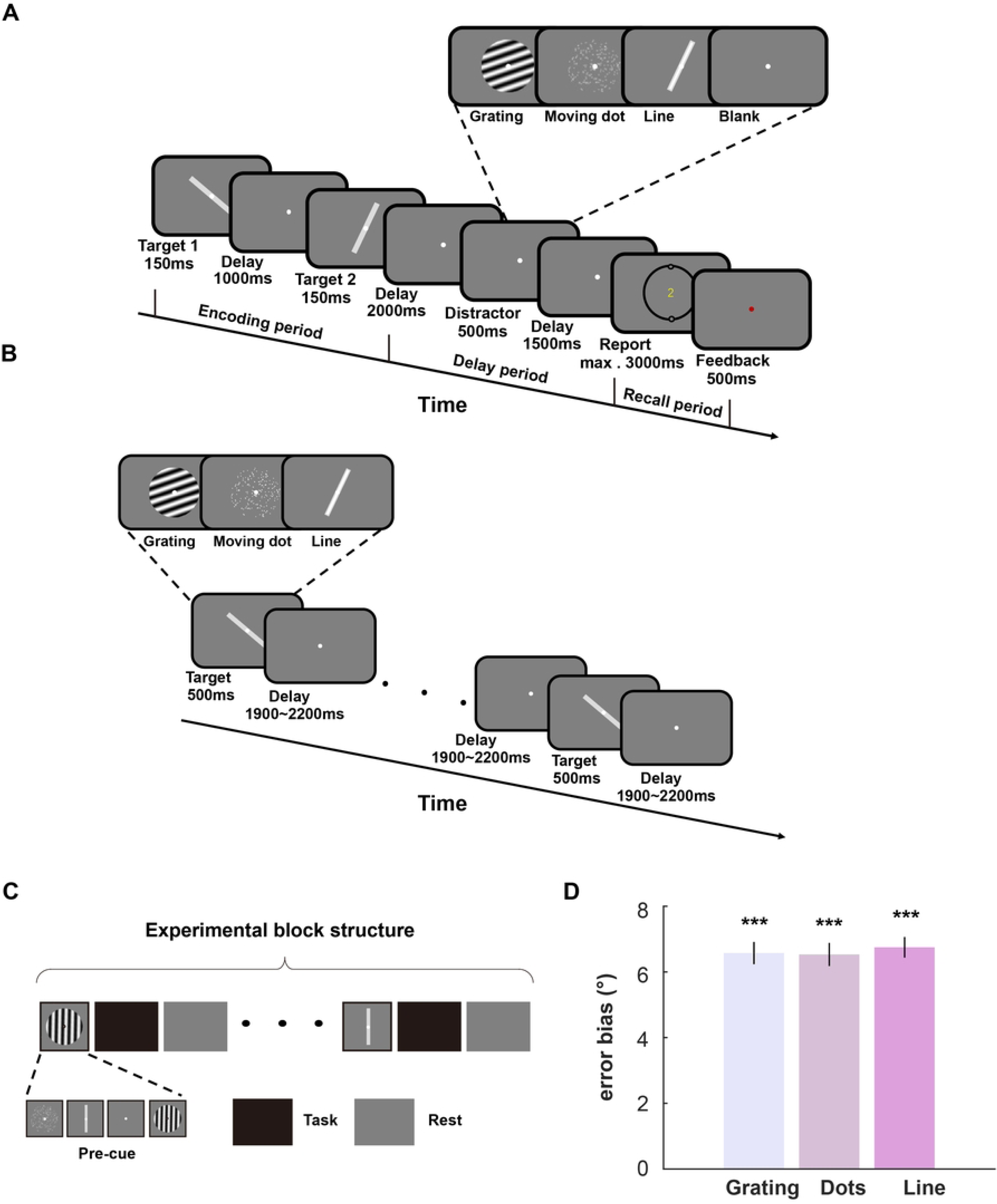
Experimental design and behavioral results. (A) VWM task. Participants sequentially viewed two oriented bars and memorized both their orientations and temporal order (first or second). During the delay, one of three distractor types, or no distractor, was presented in separate blocks. At recall, a numerical cue (“1” or “2”) indicated which bar to report, and participants adjusted a response dial to match the remembered orientation. **(B) AD task.** Stimuli identical to the distractors in the VWM task were presented in each block. Participants detected whether the stimulus orientation (for grating and line stimuli) or motion direction (for moving dots) was horizontal or vertical. **(C) Experimental block structure.** The VWM task comprised 16 blocks, each consisting of 24 trials. Each block began with a 100% valid pre-cue used to indicate the type of the block (grating, moving dot, line or blank). The AD task comprised 6 blocks, each consisting of 55 trials. Each block began with a 100% valid pre-cue used to indicate the type of the block (grating, moving dot or line). Because distractor categories in the VWM task or target categories in the AD task were grouped into separate blocks, participants could anticipate the category of the upcoming distractor or target. **(D)** Absolute values of the deviation in behavioral errors under distractor conditions compared with the condition without distractors. Error bias relative to the no-distractor condition was significant for all three distractor conditions. There were no differences in error bias between the three distractor conditions. The error bar represents ±1 SEM. \**p* < 0.05, \*\**p* < 0.01, \*\*\**p* < 0.001. The numerical data underlying panel D can be found in S1 Data.

### VWM task behavioral results

To quantify distractor-induced bias, we first calculated the absolute value of the behavioral error and, for each distractor trial, calculated the difference between its absolute error and the mean absolute error in the no-distractor condition. We then took the absolute value of this difference and averaged these values for each distractor type to obtain a single error-bias measure for each distractor condition of each participant. For example, if the mean absolute error in the no-distractor condition was 2° and the absolute error on a distractor trial was 7°, the error bias for that trial would be 5°. Similarly, if the difference was −5°, the error bias would also be 5°. Taking the absolute value allowed us to quantify the magnitude of distractor-related deviation from baseline regardless of its direction. All three distractor conditions showed significant error biases relative to the no-distractor condition (Fig 1D; grating: t(29) = 19.434, *p* < 0.001; moving dot: t(29) = 18.532, *p* < 0.001; line: t(29) = 21.532, *p* < 0.001), indicating that distractors consistently biased memory reports. A repeated-measures ANOVA showed no significant differences in error bias among the three distractor conditions (F(2,58) = 0.39; *p* = 0.679), indicating that the magnitude of distractor-induced bias was similar across distractor categories.

### Specific, persistent and stable preparation in pre-distractor brain activity

We first focused on the delay period preceding distractor onset to ask how the brain prepares for predictable task-irrelevant information before it appears. Specifically, we asked whether this preparation is distractor-specific—that is, whether different predictable distractor categories are associated with distinct anticipatory coding patterns—or whether the brain instead adopts a more generalized preparatory state. To address this question, we performed decoding analyses on the categories of the predictable distractors.

Decoding revealed significant classification performance for distractor categories before distractor onset (Fig 2A), confirming that brain activity differed across distractor categories before the distractors were physically present. This significant decoding performance (Fig 2A) emerged before the onset of the distractor, indicating that pre-distractor brain activity already carried category-specific information about the upcoming task-irrelevant stimulus. Thus, the brain did not appear to prepare for predictable distractors in a generic manner, but instead formed distinct anticipatory coding patterns for different distractor categories.

**Fig 2.**
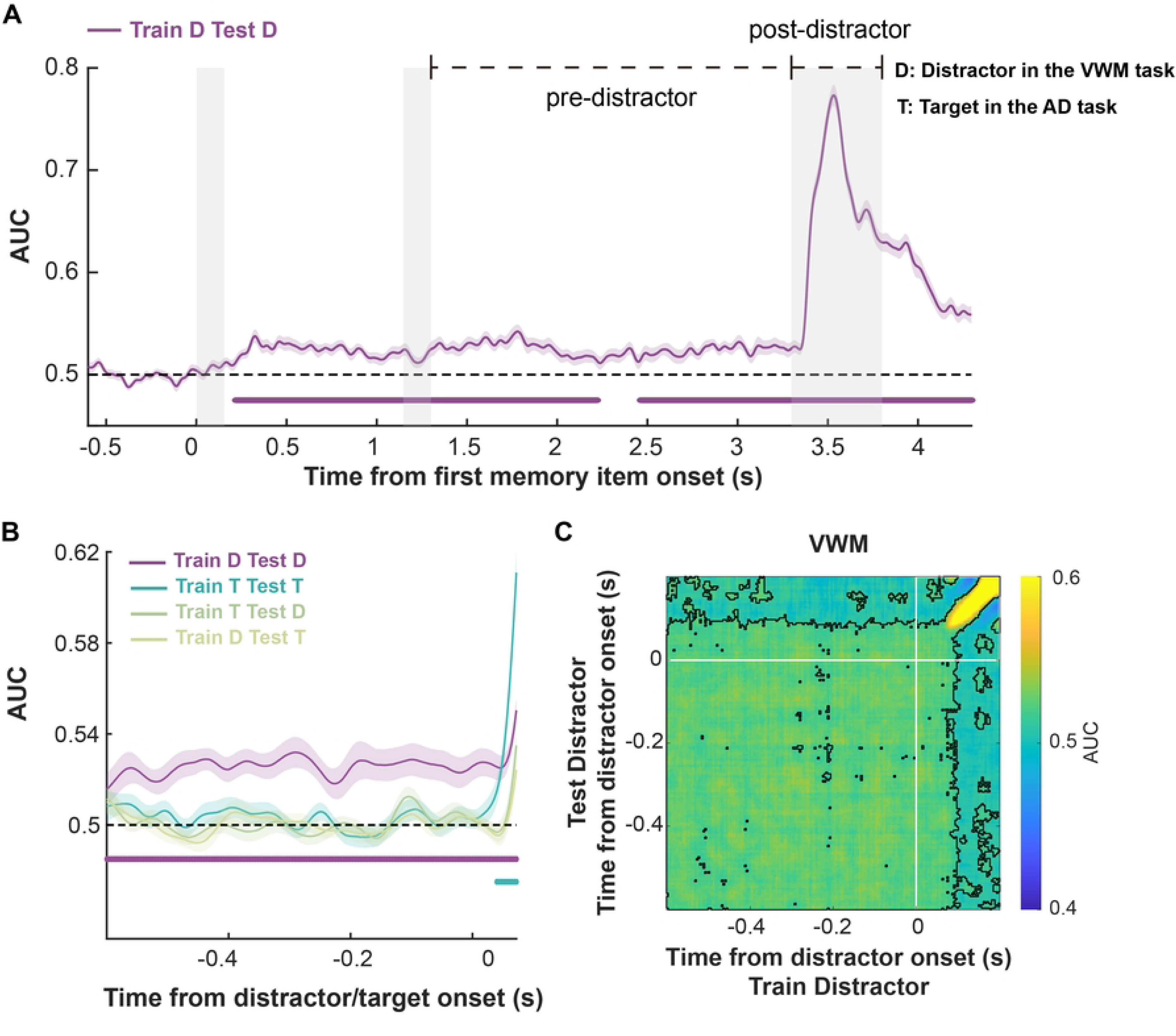
Decoding performance and temporal generalization of the distractor and target in the pre-distractor and pre-target periods. (A) Decoding time courses. Decoding performance (AUC) for the distractor category, time-locked to the onset of the first memory item. Shaded areas denote ±1 SEM. Horizontal bars indicate time clusters showing above-chance decoding (cluster-based permutation test, *p* < 0.05). Gray shading marks stimulus presentation periods (0-0.15 s for the first bar, 1.15-1.3 s for the second bar, and 3.3–3.8 s for the distractor). **(B) Decoding time courses.** Decoding performance (AUC) for each task and cross-decoding between tasks. Same analysis in (a), but time-locked to the onset of the distractor in the VWM task or the target in the AD task. Blue and purple curves show within-task decoding of the AD task target and VWM task distractor, respectively; green and yellow curves show cross-decoding performance (train on the AD target and test on the VWM distractor, and vice versa. See the color code.). Shaded areas denote ±1 SEM. Horizontal bars indicate time clusters showing above-chance decoding (cluster-based permutation test, *p* < 0.05). **(C) Temporal generalization matrices.** Train and test on distractors. Black outlines indicate matrix elements showing above-chance decoding performance (cluster-based permutation test: *p* < 0.05). Note that the lower left quadrant includes pre-distractor time points, and the upper right the post-distractor time points. The numerical data underlying panels A-C can be found in S2-S4 Data.

Because the target category in the AD task was also predictable, we next asked whether predictable task-relevant stimuli could also be decoded and, if so, whether they shared a common neural code with task-irrelevant stimuli or were represented in distinct formats. To test this, we performed within-task decoding and cross-task decoding analyses using the same stimulus categories presented as distractors during the VWM delay and as targets in the AD task. In contrast to distractor decoding, no significant above-chance cluster was observed before target onset (Fig 2B), suggesting that anticipatory coding for predictable task-irrelevant stimuli differed from that for predictable task-relevant stimuli.

We then examined the temporal structure of this pre-distractor coding using cross-temporal generalization analysis, in which a decoder was trained to discriminate distractor categories at each time point and tested across all other time points in a cross-validated manner. The temporal generalization matrix revealed significant decoding clusters that generalized across time points within the pre-distractor interval (Fig 2C, bottom left quadrant), indicating that the brain maintained a stable anticipatory coding pattern. Together, these results indicate that pre-distractor brain activity carries distractor-specific anticipatory coding that is both persistent and temporally stable.

### Flexible coding in post-distractor brain activity

We then investigated whether post-onset distractor and target representations shared a common neural code or were represented in distinct formats. We performed within-task decoding and cross-task decoding analyses using the same stimulus categories presented as distractors during the VWM delay and as targets in the AD task. Successful within-task decoding together with reduced cross-task generalization would indicate that post-distractor representations were shaped by behavioral relevance rather than maintaining a common representational format.

Figure 3A showed that both distractors in the VWM task and targets in the AD task could be decoded after stimulus onset (Fig 3A), but cross-decoding (trained on the AD targets and tested on the VWM distractors, and vice versa) between them was significantly lower than within-condition decoding (distractor: t(29) = 12.863, *p* < 0.001; target: t(29) = 13.525, *p* < 0.001). This reduction in cross-decoding emerged after distractor onset, suggesting that task-irrelevant information (distractors in the VWM task) and task-relevant information (targets in the AD task) were represented in distinct formats. Because both distractors and targets can be characterized at two representational levels—category and feature (orientation/direction)—we next examined post-distractor coding at each level in turn.

**Fig 3.**
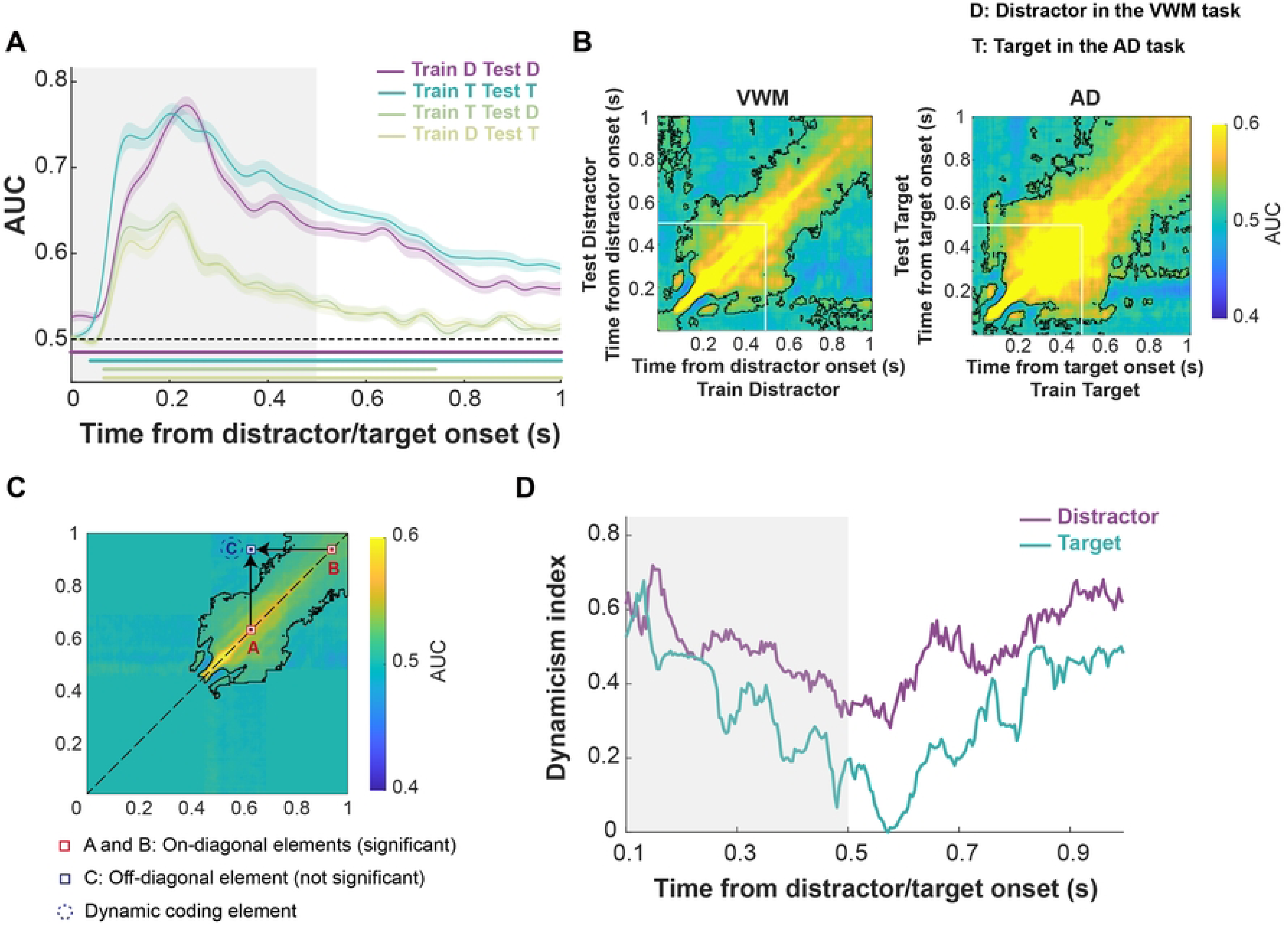
Decoding performance and temporal generalization of the distractor and target in the post-distractor and post-target periods. (A) Decoding time courses. Decoding performance (AUC) for each task and cross-decoding between tasks, time-locked to the onset of the distractor in the VWM task or the target in the AD task. Blue and purple curves show within-task decoding of the AD task target and VMW task distractor, respectively; green and yellow curves show cross-decoding performance (train on AD target and test on VWM distractor, and vice versa. See the color code). Shaded areas denote ±1 SEM. Horizontal bars indicate time clusters showing above-chance decoding (cluster-based permutation test, *p* < 0.05). Gray shading marks stimulus presentation periods (0-0.5 s for the distractor and target). **(B) Temporal generalization matrices.** Left panel: train and test on distractors in the VWM task. Right panel: train and test on targets in the AD task. Black outlines indicate matrix elements showing above-chance decoding performance (cluster-based permutation test; *p* < 0.05). **(C) Dynamic coding analysis.** Off-diagonal elements with lower decoding performance compared to both corresponding diagonal elements were defined as dynamic coding elements (see Materials and Methods for details). **(D) Dynamicism index.** Proportion of dynamic coding elements across time. High values indicate a more dynamic, non-generalizing code. Time indicated the time from 0.1 to 1 s, relative to the target (distractor) onset. Gray shading marks stimulus presentation periods. The numerical data underlying panels A, B, and D can be found in S3, S4, and S5 Data respectively.

At the category level, decoding for distractor categories after onset was more concentrated near the diagonal of the temporal generalization matrix than was target decoding (Fig 3B), indicating weaker temporal generalization and more dynamic coding. To quantify this pattern, we computed a dynamicism index (Fig 3D; see Materials and Methods), which measures the extent to which significant on-diagonal decoding fails to generalize across time. Given that decoding performance showed temporal changes approximately 0.1 s following the distractor (target) onset, we focused our analyses on the interval from 0.1 to 0.5 s following the distractor (target) onset. Using this index, we found that distractor coding was significantly more dynamic (less stable) than target coding (t(29) = 98.291, *p* < 0.001).

We next examined post-distractor coding at the feature level by decoding orientation/direction using Mahalanobis distance. VWM distractor features were significantly decoded only within a relatively brief time window beginning around 0.1 s after onset, whereas AD target features showed more sustained decoding (Fig 4A). Even within the time window in which both VWM distractor and AD target features were decodable, no significant cross-decoding was observed between them, suggesting either weaker distractor feature coding or a difference in coding format between the VWM and AD tasks.

**Fig 4.**
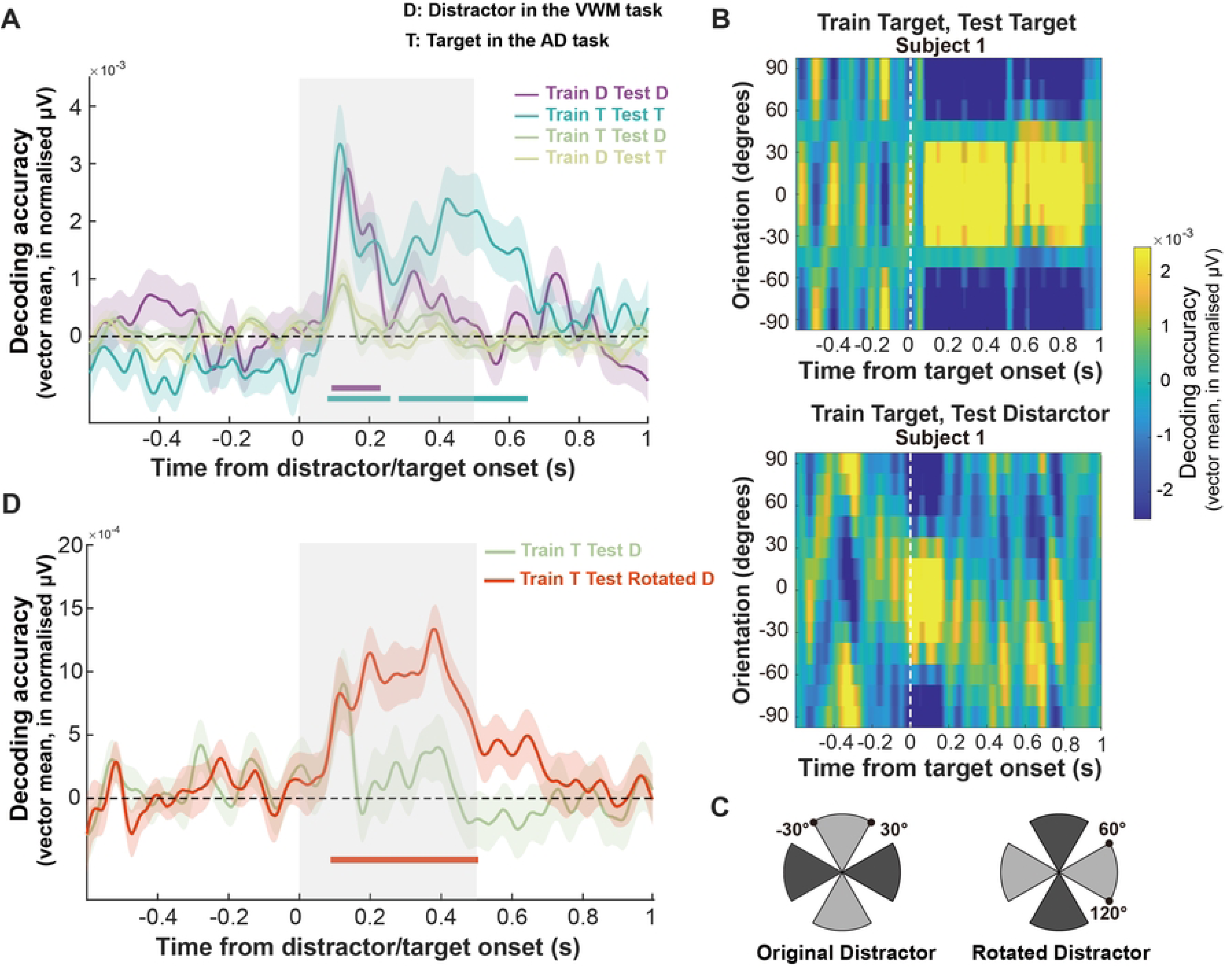
Time-resolved orientation/direction decoding. (A) Time-resolved decoding of orientation/direction. Time-resolved decoding performance derived from the vector mean of the convolved tuning curves (i.e., decoding accuracy). Horizontal bars indicate significant clusters (cluster-based permutation test, *p* < 0.05, corrected). Shades areas denote ±1 SEM. Gray shading marks the distractor or target presentation period in the VWM and AD tasks (0-0.5 s). **(B) Average decoding performance across time.** The top panel displays the time-resolved average decoding performance (mean-centered, sign-reversed Mahalanobis distance) within the AD task for target orientations/directions; the bottom panel shows the time-resolved decoding accuracy between AD and VWM tasks computed by training on the AD target and testing on the VWM distractor orientations/directions for a representative participant. **(C) Illustration of rotated coding of the distractor in the VWM task.** Mahalanobis-distance decoding trained on target orientations from the AD task and tested on either the original VWM distractor orientations (left) or on VWM distractor orientations (right) rotated in feature space (e.g., −30° rotated clockwise by 90° to 60°; see Materials and Methods). **(D) Cross-decoding of rotated distractor representations.** Same as **(A)**, but training on the AD target and testing on the original or rotated VMW distractor orientations. The numerical data underlying panels A, B and D can be found in S6-S8 Data.

To distinguish between these possibilities, we first examined the results from individual participants. Figure 4B shows data from a representative participant and provides an initial indication that distractor feature coding in the VWM task was transformed relative to target coding in the AD task. When both training and testing were performed on target features, all orientations/directions were aligned to 0° to allow averaging across trials, and the highest decoding remained centered near 0° across time (Fig 4B top). This pattern indicates maximal similarity when the trained and tested orientations/directions matched. In contrast, when training was performed on AD target features and testing on VWM distractor features, the high-decoding region was initially centered near 0° but gradually shifted over time (e.g., toward the −60° bin at ∼0.4 s), suggesting that VWM distractor representations became progressively displaced, or rotated, in feature space relative to AD target representations (Fig 4B bottom). To test this more directly at the group level, we estimated, for each participant, the rotation angle by identifying the AD target feature angle that best matched the VWM distractor representation within the 0.1–0.5 s interval (e.g., a distractor presented at 30° could be best matched by a target representation at 60°) (Fig 4C). Cross-decoding between the AD target and this best-matching rotated VWM distractor remained significantly above chance across this window (t(29) = 9.345, *p* < 0.001; Fig 4D). Together, these results suggest that post-distractor coding differs from AD target coding not only in temporal dynamicity, but also in feature-level format, consistent with a rotation of the features in the distractor representations.

### Relationship between pre-distractor and post-distractor coding in VWM

Having shown that distractor coding differed between the pre-distractor and post-distractor periods, we next asked whether these two phases of distractor processing in the VWM task were related. Specifically, we tested whether distractor-specific coding and the degree of stability before distractor onset were associated with the degree of dynamicity after distractor onset.

Because the dynamicism index was computed at the group level, we first trained classifiers separately for each participant and obtained ten estimates of decoding performance per participant. We then sampled from these estimates of all the participants to generate 1,000 random samples pairing group-level pre-distractor decoding performance and dynamicism index with the post-distractor dynamicism index and computed Pearson correlation using all the samples.

This analysis revealed the post-distractor dynamicism index was negatively correlated with both pre-distractor decoding performance (r = −0.378, *p* < 0.001, permutation *p* = 0.001; Fig 5A) and pre-distractor dynamicism index (r = −0.318, *p* < 0.001, permutation *p* = 0.001). Since a significant positive correlation was observed between pre-distractor decoding performance and the pre-distractor dynamicism index (r = 0.883, *p* < 0.001, permutation *p* = 0.001), we repeated the analysis using average pre-distractor decoding performance and dynamicism index (-2–0 s relative to distractor onset) as control variables to rule out the possibility that another variable indirectly influenced these two relationships. The partial correlation between pre-distractor decoding performance and the post-distractor dynamicism index remained significant (r = −0.219, *p* < 0.001, permutation *p* = 0.001). In contrast, the correlation between the pre-distractor dynamicism index and the post-distractor dynamicism index was not significant (r = −0.037, *p* = 0.249, permutation *p* = 0.239). This indicates that there is no independent linear relationship between the pre-distractor and post-distractor dynamicism index, suggesting that the negative correlation between the two is mediated by pre-distractor decoding performance.

**Fig 5.**
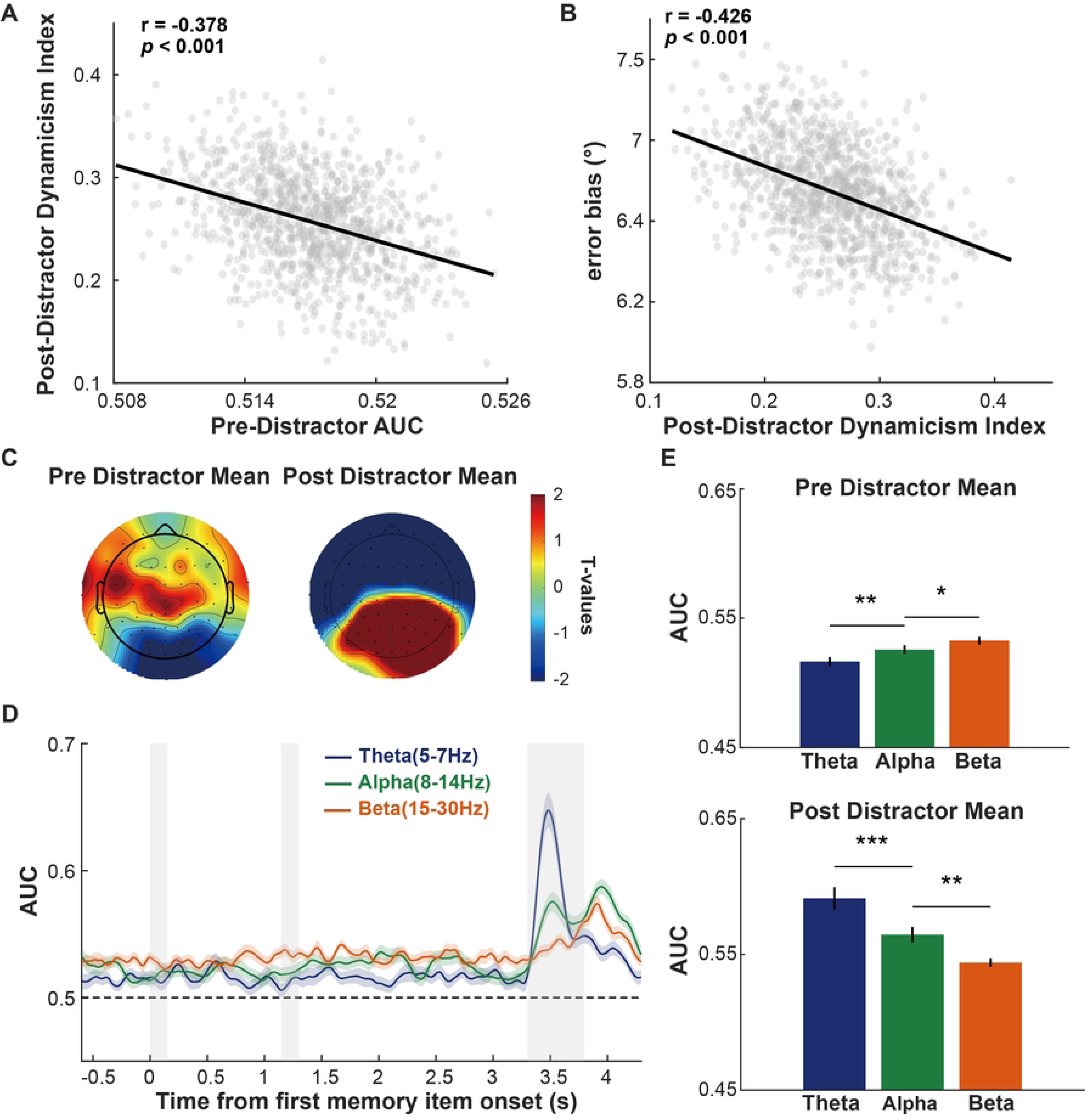
Relationship between pre-distractor coding, post-distractor dynamics, and behavioral performance. (A) Correlation between pre-distractor decoding performance and post-distractor dynamicism index. The scatter plot shows a negative relationship between mean decoding performance during the pre-distractor period (1.3-3.3 s after the first memory bar) and the mean dynamicism index during the post-distractor period (3.4–3.8 s after the first memory bar). **(B) Correlation between post-distractor dynamicism index and error bias.** The scatter plot shows a negative relationship between the mean dynamicism index during the post-distractor period (3.4–3.8 s after the first memory bar) and the mean error bias. **(C) Topographic maps of distractor categories classification weights.** T-values of the classification weights for the pre-distractor period (left) and post-distractor period (right). **(D) Decoding performance in different frequency bands.** Time-resolved decoding performance of distractor categories obtained separately for each frequency band. Memory items and the distractor were presented at 0–0.15 s, 1.15–1.3 s, and 3.3–3.8 s (gray shading). Blue: theta (5–7 Hz); green: alpha (8–14 Hz); orange: beta (15–30 Hz). **(E) Mean decoding performance during the pre-distractor and post-distractor period.** Bars depict the mean decoding performance in each of the three frequency bands during the pre-distractor (top) and post-distractor (bottom) periods. Error bars represent ±1 SEM. \**p* < 0.05, \*\**p* < 0.01, \*\*\**p* < 0.001. The numerical data underlying panels A-B can be found in S9 Data. The numerical data underlying panels C-E can be found in S10-S11 Data.

Furthermore, we examined whether distractor representations during the pre-distractor and post-distractor periods were associated with the behavioral consequences of distraction, quantified using behavioral error bias (the deviation in recall error relative to the no-distractor baseline; see Materials and Methods for details). This analysis indicated a negative correlation between the post-distractor dynamicism index and error bias (r = −0.426, *p* < 0.001, permutation *p* = 0.001; Fig 5B). Specifically, a more dynamic encoding (less stable encoding) pattern of the distractor following its presentation was associated with improved behavioral performance. When we repeated this analysis using the average pre-distractor decoding performance (-2-0 s relative to distractor onset) as the control variable, the partial correlation remained significant (r = −0.357, *p* < 0.001, permutation *p* = 0.001). Notably, a positive partial correlation was observed between pre-distractor decoding performance and error bias, controlling for the average post-distractor dynamicism index from 0.1 to 0.5 s after distractor onset (r = 0.155, *p* < 0.001, permutation *p* = 0.001). This indicates that higher pre-distractor decoding performance is associated with greater error bias.

Together, these results indicate that pre-distractor and post-distractor coding were not independent; rather, they were systematically related. The unstable coding of the distractor representation following the distractor contributed to distractor suppression.

### Dissociable neural signatures of distractor processing before and after distractor onset

A key question is whether distractor processing before and after distractor onset reflects a continuation of the same neural mechanism or is instead supported by dissociable mechanisms. We first looked into the EEG scalp topographies derived from decoding weights (t-test against 0) and found that the spatial patterns before and after distractor onset were nearly opposite (Fig 5C). This observation raised the possibility that distractor processing may be supported by different neural signatures across the two periods. To further test this possibility, we asked whether the difference between the two stages could also be detected in the spectral domain.

To do so, we decoded distractor categories separately within theta, alpha, and beta bands for each period. Distractor categories could be significantly decoded from all three frequency bands during both the pre-distractor period (theta: t(29) = 4.711, *p* < 0.001; alpha: t(29) = 7.472, *p* < 0.001; beta: t(29) = 10.738, *p* < 0.001) and the post-distractor period (theta: t(29) = 11.112, *p* < 0.001; alpha: t(29) = 11.441, *p* < 0.001; beta: t(29) = 14.128, *p* <.0001; Fig 5D).

We then compared decoding performance across frequency bands within each period. During the pre-distractor period, decoding performance increased from lower to higher frequencies (theta < alpha: t(29) = −3.105, *p* = 0.004; alpha < beta: t(29) = −2.169, *p* = 0.038; Fig 5E top). In contrast, the post-distractor period showed the opposite ordering (theta > alpha: t(29) = 4.699, *p* < 0.001; alpha > beta: t(29) = 3.631, *p* = 0.001; Fig 5E bottom). This reversed frequency profile indicates that distractor processing before and after distractor onset was dissociable not only in its temporal structure and topography, but also in the frequency ranges that best captured distractor-category information.

## Discussion

Understanding how the brain limits the impact of anticipated distractors requires examining how distractor representations unfold over time. Here, using EEG and multivariate analyses, we examined anticipated distractor representations before and after their onset in VWM. Anticipated distractor categories were represented before the actual distractors appeared, but the post-distractor coding was not a simple continuation of the anticipatory state. Instead, distractor processing was characterized by a transition from relatively stable pre-distractor coding to more dynamic post-distractor coding. These pre-and post-onset states were systematically related, yet behavioral protection was linked specifically to post-distractor dynamics. Together, these findings suggest that anticipated distractor control is organized through a temporally structured sequence of representational states.

The pre-distractor decoding results extend previous work on anticipatory distractor processing. Prior studies have shown that advance information about distractor locations, features, temporal structure, or difficulty can shape preparatory neural activity and improve selection [3, 7, 15, 16, 26, 27, 29, 33]. Here, distractors were centrally presented and did not require spatial orienting, yet upcoming task-irrelevant content could still be decoded before physical onset. This finding further reveals the content specificity of anticipatory distractor preparation: pre-distractor neural activity contained information about the specific distractor category that would subsequently appear.

The dissociation between anticipatory coding and behavioral protection helps clarify what pre-distractor coding may represent. A robust anticipatory representation does not necessarily indicate that the system has already entered an effective protective state. Instead, it may reflect an initial coding state that prepares the system for a predictable class of irrelevant input. Once the distractor appears, this initial state must be resolved against actual sensory input and the maintained memory representation. The negative relationship between anticipatory coding and post-distractor dynamics suggests that these stages are coordinated rather than independent. Stronger pre-onset coding was associated with less subsequent representational change, whereas stronger post-onset dynamics were associated with better behavioral protection. One interpretation is that stronger anticipatory preparation reduces the extent of post-onset reconfiguration required once the distractor appears. Such post-distractor dynamics may reflect the extent to which irrelevant input is reformatted, separated, or deprioritized after it enters processing, and behavioral protection depends more on how the distractor representation is reorganized after onset.

The comparison between task-relevant and task-irrelevant conditions further reveals that post-onset distractor processing involved a change in representational format. The same stimulus categories were represented differently when they served as targets in the AD task and when they served as distractors in the VWM task. This indicates that the decoded patterns did not simply reflect low-level visual properties of the stimuli. Rather, they were shaped by behavioral relevance, consistent with evidence that task goals and behavioral context can alter the format of neural representations [2, 13, 20, 37]. Feature-level biases in memory reports further showed that distractors were not completely excluded from processing: their features continued to influence maintained memory representations after onset [2, 10, 24, 25, 28]. These studies suggest that distractor control does not necessarily require eliminating irrelevant information from neural processing; instead, distractor information may remain represented while being transformed or reorganized.

The present results help specify the nature of this representational reorganization. After onset, anticipated distractor representations were rotated relative to their task-relevant format. Related work on working memory prioritization has similarly shown that information with different behavioral priorities can coexist while being maintained in distinct representational formats, supporting the idea that irrelevant information need not be eliminated to reduce interference, but may instead be functionally separated through representational reorganization [30, 32, 38, 39]. Distractor representations also became more temporally unstable after onset, and greater post-distractor instability was associated with smaller behavioral error biases, suggesting that reconfiguring irrelevant content into a less stable representational state may reduce its capacity to interfere with maintained memory. The temporal, spectral, and spatial analyses further characterized this post-onset reorganization. The transition from stable pre-distractor coding to more time-specific post-distractor coding was accompanied by changes in the spectral bands and spatial weight patterns that carried distractor information. Together, these temporal, spectral, and spatial changes indicate that post-onset distractor coding reflects a broader representational reorganization rather than a simple continuation of the anticipatory state, supporting the view that such reconfiguration may be a key process through which the impact of irrelevant information on maintained memory is reduced.

These findings also speak to dual-mechanism accounts of cognitive control. This framework distinguishes between proactive control, which prepares the system in advance, and reactive control, which is recruited after stimulus onset [4–6, 17]. Although this distinction has often been discussed in relation to task-relevant goals, cues, and responses, recent studies have begun to examine proactive and reactive periods in the suppression of task-irrelevant information [15, 26, 41]. Our results are consistent with this broader view, but add a representational account of how anticipated distractors are processed across these periods. Anticipated distractors were represented before onset, consistent with an anticipatory state, but behavioral protection was most closely associated with how their representations were reorganized after onset. Thus, task-irrelevant information may show a proactive-reactive temporal organization at the level of representational states, rather than being governed by a single preparatory mechanism.

Several limitations should be considered. First, the relationship between post-distractor dynamics and behavioral protection is correlational, so the present results do not establish a causal role for post-distractor dynamics in protecting VWM. Future studies could combine multivariate EEG with causal manipulations, such as brain stimulation or task manipulations that selectively alter post-distractor dynamics. Second, the spatial resolution of EEG limits the ability to identify the cortical sources and network interactions supporting pre-and post-distractor states. Future fMRI, MEG, or intracranial EEG studies could examine how different brain regions contribute to anticipatory coding, post-onset dynamics, and their coordination. Third, although the AD task allowed us to compare task-relevant and task-irrelevant representations for the same stimulus categories, further work is needed to determine whether similar dynamics occur for other distractor types, memory contents, and task demands.

Together, these findings clarify how anticipation and subsequent distractor processing are coordinated in distractor control. Advance coding of irrelevant content shows that the brain can prepare for upcoming distraction in a content-specific manner, but this preparatory state does not by itself determine whether VWM will be protected. Instead, protection is more closely tied to how distractor representations are reorganized after the distractor enters processing. Thus, anticipated distraction is not simply a problem of building an effective filter in advance; it is a problem of managing how irrelevant information is reformatted across the transition from expectation to sensory input. This perspective places the control of distraction in the coordinated evolution of representational states, revealing how preparation before distraction and processing after distraction are integrated within a single temporally structured mechanism.

## Materials and Methods

### Ethics statement

The study was approved by the Human Research Ethics Committee for Non-Clinical Faculties, the School of Psychology, Shenzhen University (approval number: SZU-PSY-2024-82). All participants were right-handed and provided verbal informed consent prior to participation.

### Participants

A total of 35 healthy, right-handed adults (15 females, age 18-25, s.d. = 0.4) with normal or corrected-to-normal visual acuity participated in the experiment. Five participants were excluded from analyses due to excessive EEG artifacts. Each participant completed two tasks: a VWM task and an AD task.

### Stimuli and procedure

The stimuli were generated using PsychToolBox in Matlab 2023a and presented on a 27-inch screen with a refresh rate of 240 Hz and a resolution of 2560 × 1440 pixels. Participants viewed the screen from a distance of approximately 102 cm in a darkened room. A gray background (RGB = 128, 128, 128) and a white fixation dot (0.2^°^ of visual angle in diameter) were maintained throughout the trials. Participants were instructed to fixate on this dot, except during the response period and inter-block breaks. All visual stimuli were presented at the center of the screen.

The VWM task (Fig 1A) employed a delayed-match-to-sample paradigm consisting of encoding, delay, and recall phases. Each block contained either no distractor or one type of distractor: grating distractor (4^°^ diameter, spatial frequency = 1 cycles/degree), moving dot distractor (200 white dots, each 0.11^°^ in diameter, displayed within an invisible circular area of 4^°^ diameter, moving at 5^°^ per second with 100% coherence), or line distractor (white line, 0.4^°^ wide and 4^°^ long). Each block began when participants pressed the space bar, followed by a 100% valid cue indicating the distractor type of the block. In distractor blocks, the pre-cue was the same stimulus as the distractor in the VWM task. Grating and line pre-cues were presented at 0° (vertical), and moving dots traveled at 0° (upward). In no-distractor blocks, the cue was a white fixation dot (Fig 1C). After viewing the pre-cue, participants initiated the block by pressing the space bar.

During the encoding period, participants were sequentially presented with two bars (4^°^ × 0.4^°^, RGB = 215, 215, 215), each for 0.15 s with a 1 s interval between them. Participants were instructed to memorize the orientations of the two bars as well as their order. Bar 2 was followed by a 3 s delay period. In the distractor blocks, a distractor was presented for 0.5 s, starting 2 s after the onset of the delay period. During the recall period, a retrospective cue (‘1’ or ‘2’) prompted participants to recall the orientation of the 1st or 2nd bar. At the same time, a probe dial consisting of two dots on a circle appeared vertically (at an orientation of 0^°^). The participants adjusted the dial to report the memorized orientation within 3 s and pressed the space key to submit their responses. Memory errors were computed as the angular difference between the reported and target orientations. A green fixation dot was presented for 0.5 s as feedback if the memory error was smaller than 15^°^; otherwise, a red fixation dot was shown for 0.5 s.

For each trial, the orientations of the tested bar were drawn from a uniform distribution from 20^°^ to 160^°^ in steps of 20^°^, plus a small angular jitter (±1^°^-±5^°^, randomly chosen for each trial). The tested orientations and their orders were counterbalanced across the entire experiment. The angular differences between the tested bar and the distractor were uniformly distributed across two angle differences (±60^°^ and ±80^°^). In each trial, the orientations of the non-tested bar were drawn from a uniform distribution ranging 1^°^ to 180^°^ in steps of 1^°^. In each trial, the two memory orientations were drawn independently, with a constraint that they differed at least by 30^°^. The orientations of the non-tested and distractor always differed by at least 10^°^. The orientations/directions of the distractor and the tested bar were the same across four conditions. Participants completed 16 blocks of 24 trials, with all experimental conditions counterbalanced across blocks. See Figure 1A for a trial schematic.

In addition to the VWM task, participants also completed an AD task (Fig 1B) where the targets were the same stimuli as the distractor in the VWM task. As in the VWM task a 100% valid cue indicating the stimulus type for that block: a grating, a moving-dot pattern, or a line stimulus, each block began when participants pressed the space bar (Fig 1C). The pre-cue matched the corresponding VWM distractor in stimulus. Each block contained only one type of stimulus. On each trial, a 0.5 s stimulus was presented, followed by a 1.9∼2.2 s inter-trial interval. Participants pressed a button whenever they detected that the orientation/direction was 0^°^ (vertical) or 90^°^ (horizontal) when the stimuli were lines or gratings, or that the direction was 0^°^ (upward) or 90^°^ (rightward) when the stimuli were moving dots. A response is only marked as correct if the orientation/direction is exactly 0° or 90^°^. All stimulus dimensions and orientations/directions (except for 21 stimuli at 0^°^ and 21 at 90^°^, these stimuli were not included in the subsequent analysis) were identical to those of the distractor in the VWM task. This task ensured that participants maintained attention on the stimuli and minimized motor-related contamination of neural signals associated with the stimulus orientations/directions other than 0^°^ and 90^°^. Participants completed six blocks of 55 trials, with all experimental conditions counterbalanced across blocks. See Figure 1B for a trial schematic. The order of the VWM task and the AD task were counterbalanced across participants.

### Behavioral data analyses

The analysis of behavioral error bias quantified deviations in recall error relative to the no-distractor baseline, using the average absolute error from no-distractor trials as the baseline measure. First, the average absolute error of no-distractor trials was calculated for each participant. Then this baseline error was subtracted from the absolute error of each distractor trial to quantify trial-wise error bias. Finally, we calculated the absolute value of this difference and averaged it for each distractor type to obtain a single error-bias measure for each distractor condition of each participant. A larger positive bias value indicates a greater deviation in recall error relative to baseline, reflecting stronger distractor-related effects.

### EEG acquisition

Continuous EEG data were recorded using a Brain Products BrainAmp recording system (GmbH) at a sampling rate of 1000 Hz. Sixty-four active scalp Ag/AgCI electrodes were arranged according to the international standard 10-20 system for electrode placement using a nylon head cap (EasyCap). The ground electrode was located at AFz, and all electrodes were referenced to the FCz. An additional electrode was placed near the outer canthus of the right eye to monitor eye-blink artifacts.

### EEG preprocessing

Offline EEG pre-processing was performed using EEGLAB [11] and custom MATLAB scripts. The data were downsampled to 250 Hz and high-pass filtered at 0.5 Hz to remove slow drifts. Electrical line noise was removed using the band-stop filter (48-52 Hz). Bad channels and EOG artifacts were identified by visual inspection and removed; bad channels (excluding EOG) were then interpolated from the neighboring electrodes.

In the VWM task, the data were segmented into epochs from 0.6 s before to 4.3 s after the onset of the first encoding stimulus for each condition (grating distractor, moving dot distractor, line distractor, and no-distractor). In the AD task, the data were epoched from-0.6 s to 1 s relative to the target onset for each condition (grating stimulus, moving dot stimulus, and line stimulus). The mean voltage during the pre-stimulus baseline (-0.6 to 0 s relative to stimulus onset) was subtracted from each epoch in both tasks. Data were then visually inspected on a trial-by-trial basis, and trials containing residual artifacts (e.g., excessive muscle activity and drifts) were removed. Independent component analysis (ICA) was subsequently performed to remove components corresponding to eye blinks and sustained scalp muscle activity from the data. Finally, the EEG data were re-referenced to the common average. After EEG artifact rejection, we also excluded no-response trials in the VWM task, as well as all trials at 0^°^ and 90^°^ orientations/directions and trials with incorrect responses in the AD task.

### Decoding analysis

Multivariate pattern analysis (MVPA) was applied to decode distractor categories in the VWM task and target categories in the AD task using patterns of EEG activity. Decoding of the three distractor/target categories was carried out using data from all electrodes with a multi-class support vector machine (SVM) combined with error-correcting output codes (ECOC) [12]. The ECOC model was implemented through the fitcecoc() function in MATLAB. For the VWM task, decoding was conducted at each of the 1225 time points from-0.6 s to 4.3 s relative to the onset of the first memory item. For the AD task, decoding was conducted at each of the 400 time points from-0.6 s to 1 s relative to the target onset. Training and test sets were generated independently for each time point. This analysis was conducted over pseudo-trials generated as mean averages of 2 randomly selected trials.

Decoding accuracy was computed at each time point using a fivefold cross-validation procedure. In each iteration, data randomly selected from 80% of the trials were fed to train a classifier, and the remaining 20% of the trials were used for testing. The procedure was repeated 100 times, each time with a new random partition of trials into five folds, yielding 100 decoding accuracy estimates for each time point. Decoding accuracies were then averaged across the three categories and across the 100 iterations, producing a mean decoding accuracy for each time point.

Classifier performance was quantified using the area under the receiver operating characteristic curve (AUC), computed separately for each participant. The AUC reflected the degree of separability between the stimulus categories, with 0.5 representing chance performance [8, 19]. Group-level significance was assessed by comparing mean AUC against chance (0.5) using a point-by-point two-tailed *t* test with a false discovery rate (FDR) correction for multiple comparisons [1]. The resulting decoding weights were then transformed into classification patterns that are physiologically interpretable [21].

To examine whether the distractor in the VWM task shared the same format as the target in the AD task, we used the cross-task validation approach in which the classifier was trained on one task and tested on the other. To assess the temporal stability (dynamics) of the distractor and the target representations, we also performed a temporal cross-decoding analysis. This yields a “temporal generalization matrix” showing time-by-time classification accuracy, which reveals whether the pattern of neural activity underlying classification performance remains stable or evolves dynamically over time [18]. A matrix element was defined as dynamic if the multivariate code at a given time point did not generalize to another time point. In other words, if two on-diagonal elements can be decoded significantly, yet the corresponding off-diagonal element linking them cannot be decoded significantly, the off-diagonal element was defined as a dynamic element (Fig 2D). Following previous work [10], we then computed the dynamicism index by calculating the proportion of dynamic elements across both columns and rows of the temporal generalization matrix.

To quantify the temporal dynamics of the distractor versus target in the post-distractor period, we randomly subsampled 180 trials for each participant and employed fivefold cross-validation with 10 iterations, yielding 10 decoding accuracies for each participant. We then randomly and independently sampled one decoding attempt from each participant, and repeated the procedure 30 times. We computed the dynamicism index of distractions and targets from 0.1 s to 1 s (relative to the distractor in the VWM task and the target in the AD task). The differences between the distractor and the target were then tested using t-tests on the average dynamicism index.

In a subsequent decoding analysis, the same decoding procedure was repeated across the frequency range of 3 to 30 Hz. The frequency spectrum was divided into canonical frequency bands: theta (5-7 Hz), alpha (8-14 Hz), and beta (15-30 Hz). The data for each frequency band were obtained by band-pass filtering the preprocessed EEG data for each trial using the EEGLAB eegfilt() function, followed by a Hilbert transform (e.g., analyses centered at 10 Hz were performed using a 9-11 Hz band-pass filter).

### Orientation and direction decoding

To reconstruct distractor and target orientations/directions from the EEG activity, we implemented an orientation decoding analysis based on the Mahalanobis distance [9], following the approach described in the previous work [36]. This method computes distances between the full range of possible orientations and quantifies how well the computed distances adhere to the parametric circular space of the orientation.

We performed the orientation decoding separately for each participant using all 64 channels. Trial-wise decoding rate was computed using a leave-one-trial-out cross-validation approach. At each time point, the activity pattern of a single test trial was compared to the pattern of all other trials (training trials) at the same time point. These patterns were grouped into 12 orientation bins relative to the orientations/directions of the test trial, each bin centered at-90^°^,-75^°^,-60^°^,-45^°^,-30^°^,-15^°^, 0^°^, 15^°^, 30^°^, 45^°^, 60^°^, 75^°^ (at a bin width of 30^°^).

We computed Mahalanobis distances between the test trial and each orientation bin using the covariance estimated from all training trials. A tuning curve was computed by multiplying the cosine of the center of each orientation bin (θ) with the corresponding sign-reversed distances (d(θ), higher values correspond to greater relative similarity) [31]. The vector mean of this tuning curve was taken as the decoding accuracy (da). A high value indicates that the difference between the test trial and training trials with a similar orientation/direction is smaller compared to those with different orientations/directions.

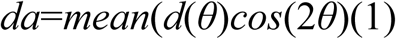

Decoding values were across all trials, and cluster-corrected sign-permutation significance tests were carried out to determine the significance of the decoding accuracy time-course.

We found that, over time, the brain’s encoding pattern for the orientation/direction of the distractor appears to shift. To test this hypothesis, in a follow-up analysis, the distractor orientations/directions were rotated by ±15^°^, ±30^°^, ± 45^°^, ±60^°^, ±75^°^ and ±90^°^, and then we compared the encoding similarity between the target orientations/directions and the rotated distractor orientations/directions. The most similar rotated distractor orientation/direction (averaged from 0.1 s to 0.5 s, relative to the distractor) to the target orientation/direction was selected for each participant to perform decoding analysis.

### Correlation analysis

To quantify the relationship between pre-distractor and post-distractor coding pattern, we performed Pearson’s correlation between the AUC and dynamicism index during the pre-distractor period (1.3 s to 3.3 s, relative to the first memory item onset) and the dynamicism index during the post-distractor period (3.4 s to 3.8 s, relative to the first memory item onset). Because the dynamicism index is a group-level measure, it cannot be directly used for the correlation analysis. To address this, we randomly subsampled 180 trials for each participant and performed decoding analysis using a fivefold cross-validation with 10 iterations. The subsampled trials used for decoding were then used to compute the dynamicism index at the group-level. This procedure was repeated 1,000 times, producing 1,000 pairs of group-level AUC and dynamicism index for correlation. To assess the significance of each correlation, we performed a permutation test. One variable was held fixed while the paired values of the other variable were randomly shuffled across the 1,000 data points to disrupt their correspondence. This shuffling procedure was repeated 1,000 times, and the correlation coefficient was recalculated for each permutation to generate a null distribution. The significance of the empirical correlation coefficient was determined by its position within this null distribution using a two-tailed test. This procedure was used for all correlation analyses reported in this study.

## Data availability statement

The individual numerical data underlying the main figures are provided in the Supporting Information files (S1–S11 Data). All analysis code and raw data are publicly available on Zenodo(https://doi.org/10.5281/zenodo.22677862).

## Abbreviations

EEG: electroencephalography
MVPA: Multivariate Pattern Analysis
VWM: visual working memory
AD: angle detection
AUC: area under the receiver operating characteristic curve.

## Supporting information

S1 Data. Underlying numerical data for graphs in Fig 1D.

S2 Data. Underlying numerical data for graphs in Fig 2A.

S3 Data. Underlying numerical data for graphs in Fig 2B and 3A.

S4 Data. Underlying numerical data for graphs in Fig 2C and 3B.

S5 Data. Underlying numerical data for graphs in Fig 3D.

S6 Data. Underlying numerical data for graphs in Fig 4A.

S7 Data. Underlying numerical data for graphs in Fig 4B.

S8 Data. Underlying numerical data for graphs in Fig 4D.

S9 Data. Underlying numerical data for graphs in Fig 5A and 5B.

S10 Data. Underlying numerical data for graphs in Fig 5C.

S11 Data. Underlying numerical data for graphs in Fig 5D and 5E.

## Notes

### Competing Interest Statement

The authors have declared no competing interest.

## References

1. Benjamini Y, Hochberg Y. Controlling the false discovery rate: a practical and powerful approach to multiple testing. Journal of the Royal statistical society: series B (Methodological). 1995;57(1):289–300.

2. Bettencourt KC, Xu Y. Decoding the content of visual short-term memory under distraction in occipital and parietal areas. Nature neuroscience. 2016;19(1):150–7.

3. Bonnefond M, Jensen O. Alpha oscillations serve to protect working memory maintenance against anticipated distracters. Current biology. 2012;22(20):1969–74.

4. Braver TS. The variable nature of cognitive control: a dual mechanisms framework. Trends in cognitive sciences. 2012;16(2):106–13.

5. Braver TS, Gray JR, Burgess GC. Explaining the many varieties of working memory variation: Dual mechanisms of cognitive control. Variation in working memory. 2007;75(106):10.1093.

6. Braver TS, Paxton JL, Locke HS, Barch DM. Flexible neural mechanisms of cognitive control within human prefrontal cortex. Proceedings of the National Academy of Sciences. 2009;106(18):7351–6.

7. Chelazzi L, Marini F, Pascucci D, Turatto M. Getting rid of visual distractors: The why, when, how, and where. Current opinion in psychology. 2019;29:135–47.

8. Chen Y-T, van Ede F, Kuo B-C. Alpha oscillations track content-specific working memory capacity. Journal of Neuroscience. 2022;42(38):7285–93.

9. De Maesschalck R, Jouan-Rimbaud D, Massart DL. The mahalanobis distance. Chemometrics and intelligent laboratory systems. 2000;50(1):1–18.

10. Degutis JK, Weber S, Soch J, Haynes J-D. Neural dynamics of visual working memory representation during sensory distraction. Elife. 2025;13:RP99290.

11. Delorme A, Makeig S. EEGLAB: an open source toolbox for analysis of single-trial EEG dynamics including independent component analysis. Journal of neuroscience methods. 2004;134(1):9–21.

12. Dietterich TG, Bakiri G. Solving multiclass learning problems via error-correcting output codes. Journal of artificial intelligence research. 1994;2:263–86.

13. Duncan J. An adaptive coding model of neural function in prefrontal cortex. Nature reviews neuroscience. 2001;2(11):820–9.

14. Failing M, Wang B, Theeuwes J. Spatial suppression due to statistical regularities is driven by distractor suppression not by target activation. Attention, Perception, & Psychophysics. 2019;81(5):1405–14.

15. Ferrante O, Jensen O, Hickey C. Predictive distractor processing relies on integrated proactive and reactive attentional mechanisms. Journal of Neuroscience. 2026.

16. Gaspelin N, Luck SJ. The role of inhibition in avoiding distraction by salient stimuli. Trends in cognitive sciences. 2018;22(1):79–92.

17. Gonthier C, Braver TS, Bugg JM. Dissociating proactive and reactive control in the Stroop task. Memory & Cognition. 2016;44(5):778–88.

18. Grootswagers T, Wardle SG, Carlson TA. Decoding dynamic brain patterns from evoked responses: a tutorial on multivariate pattern analysis applied to time series neuroimaging data. Journal of cognitive neuroscience. 2017;29(4):677–97.

19. Hand DJ, Till RJ. A simple generalisation of the area under the ROC curve for multiple class classification problems. Machine learning. 2001;45(2):171–86.

20. Harel A, Kravitz DJ, Baker CI. Task context impacts visual object processing differentially across the cortex. Proceedings of the National Academy of Sciences. 2014;111(10):E962–E71.

21. Haufe S, Meinecke F, Görgen K, Dähne S, Haynes J-D, Blankertz B, et al. On the interpretation of weight vectors of linear models in multivariate neuroimaging. Neuroimage. 2014;87:96–110.

22. Jensen O, Mazaheri A. Shaping functional architecture by oscillatory alpha activity: gating by inhibition. Frontiers in human neuroscience. 2010;4:186.

23. Liesefeld HR, Liesefeld AM, Sauseng P, Jacob SN, Müller HJ. How visual working memory handles distraction: Cognitive mechanisms and electrophysiological correlates. Visual Cognition. 2020;28(5-8):372–87.

24. Lorenc ES, Mallett R, Lewis-Peacock JA. Distraction in visual working memory: Resistance is not futile. Trends in cognitive sciences. 2021;25(3):228–39.

25. Lorenc ES, Sreenivasan KK, Nee DE, Vandenbroucke AR, D’Esposito M. Flexible coding of visual working memory representations during distraction. Journal of Neuroscience. 2018;38(23):5267–76.

26. Marini F, Demeter E, Roberts KC, Chelazzi L, Woldorff MG. Orchestrating proactive and reactive mechanisms for filtering distracting information: Brain-behavior relationships revealed by a mixed-design fMRI study. Journal of Neuroscience. 2016;36(3):988–1000.

27. Noonan MP, Adamian N, Pike A, Printzlau F, Crittenden BM, Stokes MG. Distinct mechanisms for distractor suppression and target facilitation. Journal of Neuroscience. 2016;36(6):1797–807.

28. Rademaker RL, Chunharas C, Serences JT. Coexisting representations of sensory and mnemonic information in human visual cortex. Nature neuroscience. 2019;22(8):1336–44.

29. Richter D, van Moorselaar D, Theeuwes J. Proactive distractor suppression in early visual cortex. Elife. 2025;13:RP101733.

30. Ritz H, Shenhav A. Orthogonal neural encoding of targets and distractors supports multivariate cognitive control. Nature Human Behaviour. 2024;8(5):945–61.

31. Van Ede F, Chekroud SR, Stokes MG, Nobre AC. Decoding the influence of anticipatory states on visual perception in the presence of temporal distractors. Nature communications. 2018;9(1):1449.

32. van Loon AM, Olmos-Solis K, Fahrenfort JJ, Olivers CN. Current and future goals are represented in opposite patterns in object-selective cortex. elife. 2018;7:e38677.

33. van Moorselaar D, Lampers E, Cordesius E, Slagter HA. Neural mechanisms underlying expectation-dependent inhibition of distracting information. elife. 2020;9:e61048.

34. Wang B, Theeuwes J. How to inhibit a distractor location? Statistical learning versus active, top-down suppression. Attention, Perception, & Psychophysics. 2018;80(4):860–70.

35. Wang B, Theeuwes J. Statistical regularities modulate attentional capture. Journal of Experimental Psychology: Human Perception and Performance. 2018;44(1):13.

36. Wolff MJ, Jochim J, Akyürek EG, Stokes MG. Dynamic hidden states underlying working-memory-guided behavior. Nature neuroscience. 2017;20(6):864–71.

37. Woolgar A, Hampshire A, Thompson R, Duncan J. Adaptive coding of task-relevant information in human frontoparietal cortex. Journal of Neuroscience. 2011;31(41):14592–9.

38. Xu Y. The human posterior parietal cortices orthogonalize the representation of different streams of information concurrently coded in visual working memory. PLoS Biology. 2024;22(11):e3002915.

39. Yu Q, Teng C, Postle BR. Different states of priority recruit different neural representations in visual working memory. PLoS biology. 2020;18(6):e3000769.

40. Zhao C, Kong Y, Li D, Huang J, Kong L, Li X, et al. Suppression of distracting inputs by visual-spatial cues is driven by anticipatory alpha activity. PLoS biology. 2023;21(3):e3002014.

41. Zhao G, Chen J, Duan Y, Li S, Wang Q, Li D. The proactive and reactive mechanisms of learned spatial suppression. Cerebral Cortex. 2024;34(8):bhae333.

